# SARS-CoV-2 preS dTM vaccine booster candidates increase functional antibody responses and cross-neutralization against SARS-CoV-2 variants of concern in non-human primates

**DOI:** 10.1101/2021.09.20.461023

**Authors:** Vincent Pavot, Catherine Berry, Michael Kishko, Natalie G. Anosova, Dean Huang, Tim Tibbitts, Alice Raillard, Sylviane Gautheron, Cindy Gutzeit, Marguerite Koutsoukos, Roman Chicz, Valerie Lecouturier

**Affiliations:** Sanofi Pasteur, Marcy l’Etoile, France; Sanofi Pasteur, Cambridge, MA, USA; GSK, Rixensart, Belgium; GSK, Wavre, Belgium

## Abstract

The emergence of severe acute respiratory syndrome coronavirus 2 (SARS-CoV-2) variants that partly evade neutralizing antibodies has raised concerns of reduced vaccine effectiveness and increased infection. We previously demonstrated in preclinical models and in human clinical trials that our SARS-CoV-2 recombinant spike protein vaccine adjuvanted with AS03 (CoV2 preS dTM-AS03) elicits robust neutralizing antibody responses in naïve subjects. Here, the objective was to document the potency of various booster vaccine formulations in macaques previously vaccinated with mRNA or protein subunit vaccine candidates.

We show that one booster dose of AS03-adjuvanted CoV2 preS dTM, D614 (parental) or B.1.351 (Beta), in monovalent or bivalent (D614 + B.1.351) formulations, significantly boosted pre-existing neutralizing antibodies and elicited high and stable cross-neutralizing antibodies covering the four known SARS-CoV-2 variants of concern (Alpha, Beta, Gamma and Delta) and, unexpectedly, SARS-CoV-1, in primed macaques. Interestingly, the non-adjuvanted CoV2 preS dTM B.1.351 vaccine formulation also significantly boosted and broadened the neutralizing antibody responses.

Our findings show that these vaccine candidates used as a booster have the potential to offer cross-protection against a broad spectrum of variants. This has important implications for vaccine control of SARS-CoV-2 variants of concern and informs on the benefit of a booster with our vaccine candidates currently under evaluation in phase 2 and 3 clinical trials (NCT04762680 and NCT04904549).

## Introduction

Emergence of severe acute respiratory syndrome coronavirus 2 (SARS-CoV-2) variants of concern (VoC), Alpha in late 2020 and Delta in spring 2021, have sparked concerns about transmissibility and pathogenicity^1, 2, 3^. These VoC harbor specific mutations in the spike protein impacting transmission and antigenicity reflecting growing immune pressure at positions shown to involve antibody recognition^4^. Some VoC (Beta, Gamma and Delta) have also demonstrated partial evasion of natural and vaccine-elicited neutralizing antibodies^5, 6, 7, 8^ and are associated with reduction of vaccine effectiveness^9, 10, 11^. In response to the rapid evolution of SARS-CoV-2 and the global circulation of VoC since the end of 2020, vaccines using a spike antigen modified to contain the mutations identified in the variants are being tested in a clinical trial (NCT04785144). This strategy is based on the influenza vaccine model where the vaccine is modified yearly according to the epidemiology, likewise, a combination of spike antigens covering multiple co-circulating strains may be necessary. In early 2021, we and others selected the Beta spike (B.1.351) as it displayed the greatest breakthrough infections against the parental (D614) vaccines^9, 10^. Considering the risk of waning immunity after natural infection or immunization and the risk of antibody escape by emerging variants, the key attributes of future vaccines will be the ability to boost, prolong, and broaden protective immunity against COVID-19 and new emerging variants.

Here we formulated soluble prefusion-stabilized spike trimers (CoV2 preS dTM) with the well characterized adjuvant AS03, an oil-in-water emulsion composed of α-tocopherol, squalene and polysorbate 80^12^. AS03 potently induces antibody responses and has been shown to increase vaccine durability, promote heterologous strain cross-reactivity and to have dose-sparing effects^13, 14^. We previously reported that CoV2 preS dTM-AS03 (D614) confers protective efficacy against SARS-CoV-2 (USA-WA1/2020 isolate) challenge in non-human primates and that vaccine-induced IgG mediated protection from SARS-CoV-2 challenge following passive transfer to hamsters^15^. We also showed safety and immunogenicity in humans in a phase 1/2 clinical trial^16^ and, an optimized vaccine formulation is being assessed in a dose-ranging phase 2 clinical trial (NCT04762680) and a phase 3 efficacy trial (NCT04904549).

In this work, the objective was to assess a subunit vaccine booster in macaques vaccinated 7 months previously with either mRNA-LNP or subunit CoV2 preS dTM-AS03 Sanofi Pasteur vaccine candidates. Various formulations, AS03-adjuvanted parental (D614), variant (B.1.351), bivalent (D614+B.1.351) CoV2 preS dTM or non-adjuvanted CoV2 preS dTM (B.1.351), were evaluated for their ability to induce cross-neutralization against known VoC. Using a lentivirus-based pseudovirus (PsV) neutralizing antibody (NAb) assay^17, 18^ we assessed NAb responses against the SARS-CoV-2 D614G strain (which replaced in circulation the parental D614 strain), the four currently described VoC: Alpha (B.1.1.7), Beta (B.1.351), Gamma (B.1.1.28/P.1) and Delta (B.1.617.2) and the SARS-CoV-1 to further explore the breadth of neutralization.

## Results

### One booster dose of CoV2 preS dTM boosts pre-existing neutralizing antibodies against D614G with significant AS03-adjuvant effect

Following primary vaccination with either mRNA-LNP or CoV2 preS dTM-AS03, we assessed the neutralizing antibody (NAb) responses against the D614G. Using a GFP-based pseudovirus NAb assay, the mean NAb titers at day 35 (D35; 2 weeks post-dose 2) against the D614G pseudovirus were 2.6 log_10_ for the mRNA-primed cohort and 2.9 log_10_ for the subunit-primed cohort (Fig. 1b). Six months post-dose 2 (baseline pre-boost), mean D614G NAb titers declined to 1.4 log_10_ for the mRNA-primed cohort and to 2.2 log_10_ for the subunit-primed cohort. D614G NAb titers post-prime and at baseline pre-boost were lower in the mRNA-primed cohort compared to the subunit cohort. For comparison, the mean D614 NAb titer in a panel of 93 human convalescent sera (collected within 3 months after positive PCR test) was 2.0 log_10_ (D614 and D614G NAb titers were shown to be similar in a concordance analysis, data not shown) and the WHO International Standard for anti-SARS-CoV-2 immunoglobulin (human) (NIBSC code: 20/136) had a titer of 2.8 log_10_.

**Fig.1.**
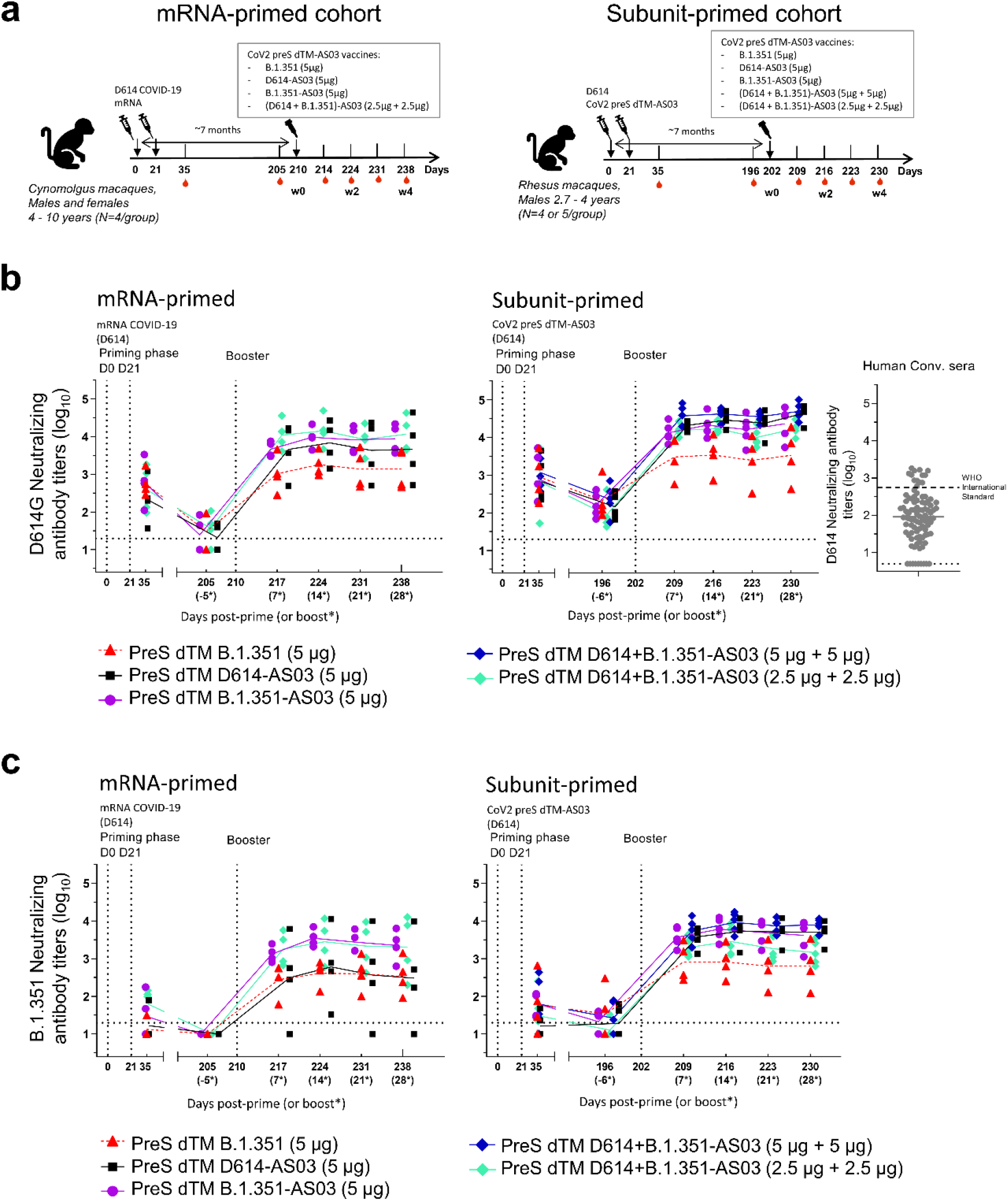
Booster neutralizing antibody responses (D614G and B.1.351/Beta) in mRNA- or subunit-primed macaques. **a** Schematic representation of the study schedule. In mRNA-primed cohort, 4 groups of 4 cynomolgus macaques were immunized intramuscularly with mRNA COVID-19 (D614) vaccine candidates on day 0 (D0) and on day 21 (D21). In subunit-primed cohort, 5 groups of 4 to 5 rhesus macaques were immunized intramuscularly with CoV2 preS dTM-AS03 (D614) vaccine candidates on D0 and D21. Both cohorts were boosted 7 months post-dose 1 with non-adjuvanted monovalent (B.1.351) CoV2 preS dTM or AS03-adjuvanted monovalent (D614 or B.1.351) or bivalent (D614+B.1.351) CoV2 preS dTM. Humoral immune responses were assessed at different timepoints throughout the study. **b** Pseudovirus neutralizing antibody (NAb) titers against the SARS-CoV-2 D614G and **c** B.1.351 (Beta) variant were assessed at different timepoints after the priming phase and after the booster immunization in macaques. Individual macaque data are shown. Connecting lines indicate mean responses and horizontal dotted lines the limits of quantification of the assay. Asterisks = timepoints relative to boosters. A human convalescent panel (N=93) was assayed against D614 and is plotted as a comparator. Dotted line represents the WHO International Standard for anti-SARS-CoV-2 immunoglobulin (human) NIBSC code: 20/136.

A booster dose with non-adjuvanted CoV2 preS dTM (B.1.351) significantly increased D614G NAb titers compared to baseline as soon as 7 days after injection, reaching a plateau in 14 days in both mRNA-primed (mean of 3.3 log_10_) and subunit-primed (mean of 3.5 log_10_) cohorts (p-values<0.001). As in each cohort, macaques were randomized for their baseline titers 6 months post-primary vaccination and peak titers post-prime (D35), we analyzed the fold-increase of NAb titers 14 days post-booster compared to baseline or post-prime. Mean increase in NAbs from baseline were 60-fold (p-value<0.001) in mRNA-primed and 16-fold (p-value<0.001) in subunit-primed cohorts. Of note, baseline titers in the mRNA cohort were lower than in the subunit cohort. The mean D614G NAb titers 14 days post-booster were significantly higher than the peak titer post-priming vaccination at D35 (3.2-fold, p-value<0.05 and 3.8-fold, p-value<0.01 for mRNA-primed and subunit-primed cohorts respectively).

A booster dose with adjuvanted CoV2 preS dTM-AS03 vaccine candidates (monovalent or bivalent formulations) also significantly increased D614G NAb titers compared to baseline in both cohorts as early as 7 days after injection and by 14 days the mean titers were 4.0 log_10_ in mRNA-primed (mean fold-increase of 370 from baseline, p-values<0.001) and 4.4 log_10_ in subunit-primed (mean fold-increase of 177 from baseline, p-values<0.001) animals.

In both the mRNA- and subunit-primed cohorts, after the AS03-adjuvanted booster, the D614G NAb titers (at all timepoints post-boost) were significantly higher than the peak titers post-priming vaccination at D35 (mean fold-increase of 21 in the mRNA-primed cohort and a mean fold-increase of 31 in the subunit-primed cohort; p-values<0.001). There were no statistically significant differences in D614G NAb titer increase (from D35 post-primary vaccination or from baseline pre-booster) between the various AS03-adjuvanted vaccine formulations.

In both cohorts, a significant adjuvant effect was observed on the fold-increase from baseline (D205 and D196) to 2 weeks post-boost with CoV2 preS dTM-AS03 (B.1.351) vaccine compared to non-adjuvanted CoV2 preS dTM (B.1.351). The increase with AS03 was 6.5-fold, (p-value<0.01) in the mRNA-primed cohort, and 8.1-fold (p-value<0.001) in the subunit-primed cohort.

The D614G NAb titers elicited by non-adjuvanted CoV2 preS dTM (B.1.351) booster were all higher than the mean D614G NAb titers of human convalescent sera (2.2 Log_10_), and all AS03-adjuvanted boost candidate vaccines induced titers above the highest titer observed in human convalescent sera in the subunit-primed cohort (the lowest D614G NAb titers was 3.6 log_10_ after the AS03-adjuvanted boost, and the highest D614G NAb titer in human convalescent sera was 3.3 log_10_).

### One booster dose of CoV2 preS dTM significantly increases Beta variant (B.1.351) neutralizing antibodies with significant AS03-adjuvant effect

We next assessed NAb titers against the Beta variant PsV (B.1.351) using the lentivirus-based neutralization assay. After the primary immunization regimen of D0 and D21, NAb titers against Beta variant were low to undetectable in the mRNA-primed cohort (mean titer of 1.4 log_10_) and low in the subunit-primed cohort (mean titer of 1.6 log_10_) at D35 (Fig. 1c). Six months post-dose 2 (D205 and D196), B.1.351 NAb titers were undetectable in the mRNA-primed cohort (<LOQ) and low to undetectable in the subunit-primed cohort (mean titer of 1.3 log_10_).

Booster dose with the non-adjuvanted CoV2 preS dTM (B.1.351) vaccine induced NAb titers against the Beta variant within 7 days after injection, with a mean titer of 2.6 log_10_ in the mRNA-primed and 2.9 log_10_ in the subunit-primed cohorts 14 days post-injection that were stable until D28. The increase in NAb titers against the Beta variant from baseline (D205 and D196) were higher with the AS03-adjuvanted CoV2 preS dTM (B.1.351) vaccine candidate compared to non-adjuvanted CoV2 preS dTM (B.1.351) in the mRNA-primed cohorts (8.5-fold, p=0.057) and significantly higher in the subunit-primed cohort (11.8-fold, p-value<0.001). No statistical differences were observed between the various AS03-adjuvanted vaccine formulations with B.1.351 mean NAb titers of 3.3 log_10_ in the mRNA-primed animals (mean fold-increase of 180 from baseline) and 3.7 log_10_ in the subunit-primed animals (mean fold-increase of 280 from baseline) 14 days post-injection (D224 and D216 respectively). In the two cohorts, after the AS03-adjuvanted booster immunization, the mean B.1.351 NAb titers observed at all time points collectively were significantly higher than the peak titers post-primary vaccination on D35 whatever the vaccine formulations (p-values<0.001, average fold-increase of 46 in the mRNA-primed and 140 in the subunit-primed cohort).

### One booster dose of CoV2 preS dTM significantly increases cross-neutralizing antibodies against the four VoC (Alpha, Beta, Gamma, Delta) and SARS-CoV-1 with significant AS03-adjuvant effect

The breadth of neutralization was further assessed against the other known SARS-CoV-2 VoC, Alpha (B.1.1.7), Gamma (B.1.1.28/P1), Delta (B.1.617.2), and the first SARS-CoV-1 from 2002/2003. NAb cross-reactivity against the three VoC was observed 14 days post-injection with all vaccine formulations in both cohorts (Fig. 2 and supp Fig. 1). The AS03-adjuvanted vaccine boosters induced, in the mRNA-primed cohort, mean VoC NAb titers of 4.0 log_10_ (Alpha), 3.6 log_10_ (Gamma) and 3.7 log_10_ (Delta) (Fig. 2a) and, in the subunit-primed cohort, mean VoC NAb titers of 4.4 log_10_ (Alpha), 4.0 log_10_ (Gamma) and 4.1 log_10_ (Delta) (Fig. 2b). As observed with the D614G and the B.1.351 variants, a significant adjuvant effect was observed for VoC B.1.1.7, B.1.1.28/P1 and B.1.617.2 NAb titer, that increased from baseline with the CoV2 preS dTM B.1.351 monovalent vaccine booster in both cohorts (mean fold-increase: 7.4, 11.9 and 7.1 for B.1.1.7, B.1.1.28/P1 and B.1.617.2 respectively in the mRNA-primed cohort, p-values<0.05 and 5.7, 8.6 and 6.6 for B.1.1.7, B.1.1.28/P1 and B.1.617.2 respectively in the subunit-primed cohort, p-values<0.05).

**Fig. 2.**
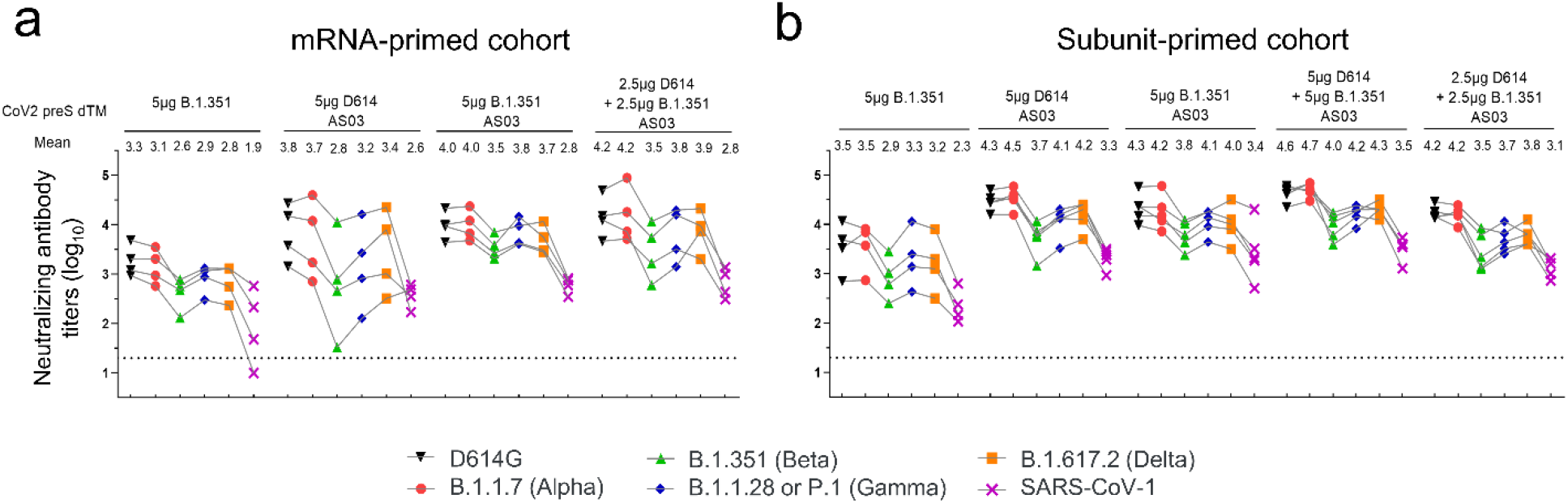
Booster cross-neutralizing antibody responses against the current VoC and SARS-CoV-1 in the mRNA- and subunit-primed macaques. Pseudovirus neutralizing antibody (NAb) assays against the SARS-CoV-2 D614G, variants of concerns (Alpha, Beta, Gamma and Delta) and SARS-CoV-1 were assessed at day 14 after the booster injection in primed-macaques **(a)** mRNA cohort and **(b)** subunit cohort. Individual macaque data are shown and are linked by connecting lines. Means are depicted above graphs. Horizontal dotted lines indicate the limits of quantification of the assay.

Unexpectedly, we also observed a significant enhanced neutralizing breadth with induction of NAb against SARS-CoV-1 14 days post-injection with all booster vaccine formulations (p-values<0.001). The non-adjuvanted vaccine booster induced significant SARS-CoV-1 neutralizing titers in all macaques except one (mean NAb titers of 1.9 log_10_ and 2.3 log_10_ in the mRNA-primed and subunit-primed cohorts respectively, p-values<0.001). AS03-adjuvanted vaccine formulations induced significant SARS-CoV-1 NAb in all macaques, with mean titers of 2.7 log_10_ and 3.3 log_10_ in the mRNA-primed cohort and subunit-primed cohort respectively, when all AS03-vaccine formulations were pooled (p-values<0.001; Fig. 2 and Supp Fig. 2). The AS03 adjuvant effect was confirmed on SARS-CoV-1 NAb titer increases from baseline when comparing the AS03-adjuvanted and the non-adjuvanted B.1.351 monovalent vaccine candidates (9.1-fold and 7-fold higher booster effect for mRNA-primed and subunit-primed cohort respectively, p-values <0.01).

### One booster dose of CoV2 preS dTM specifically increases functional antibodies

To better assess the impact of the booster on the quality of antibody responses, we calculated the individual ratio between D614G neutralizing and S-binding antibodies using serum collected at D35 and 14 days post-booster injection (D224 and D216) (Fig. 3). The ratios were significantly increased in all groups of the subunit-primed cohort from 3.1 to 7.1-fold post-booster (p-values<0.01). Although the same trend was observed in the mRNA-primed cohort, the effect was significant only in the bivalent AS03-adjuvanted booster group (6.3-fold, p-value<0.05) indicating a higher proportion of neutralizing antibodies among the S-specific antibodies. The ratios were higher than those calculated in the 93 human convalescent sera.

**Fig. 3.**
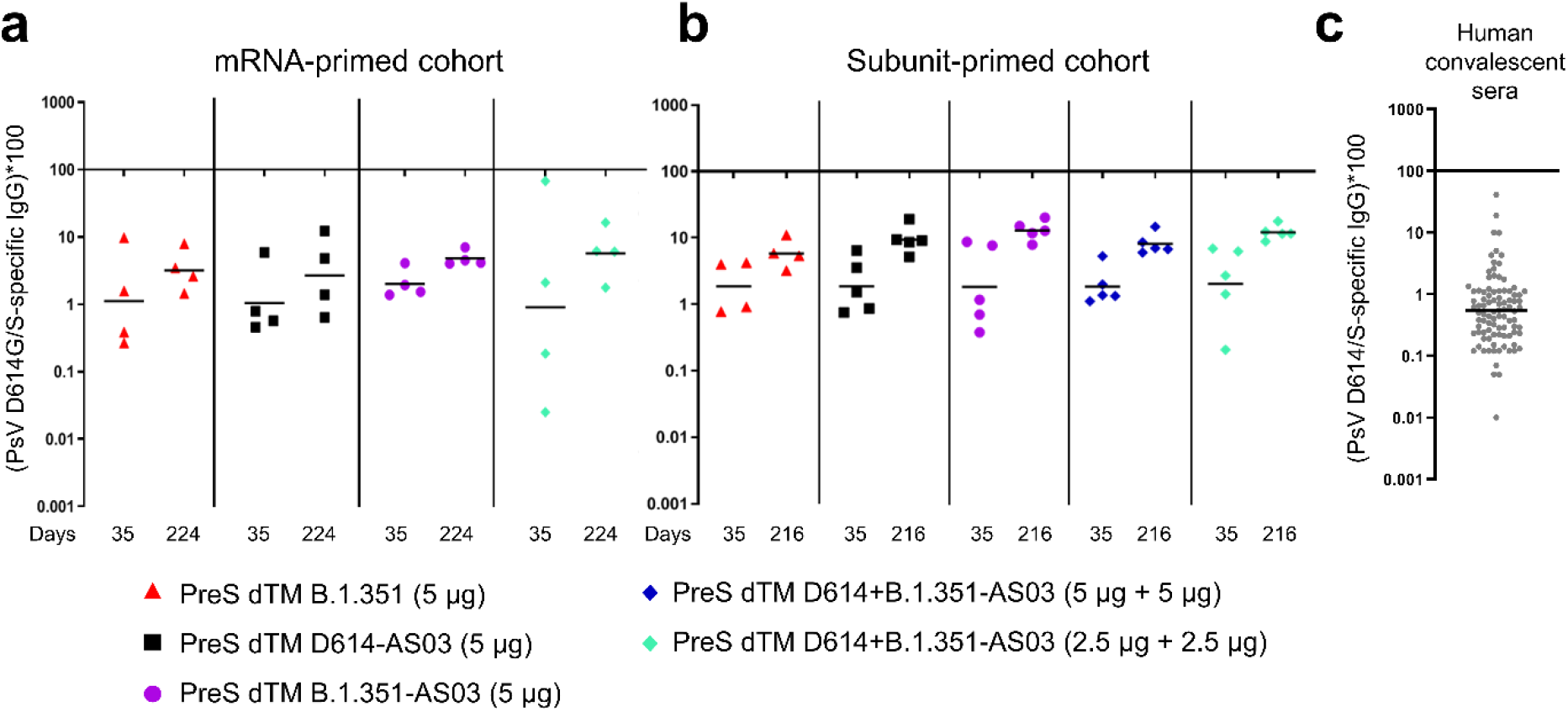
Booster immunization elicits a higher quality of antibody response in mRNA- and subunit-primed macaques. Pseudovirus D614G NAb / S-binding IgG ratio post-prime (D35) and 2 weeks post-boost (D224 or D216) for **(a)** mRNA-primed cohort and for **(b)** subunit-primed cohort and **(c)** pseudovirus D614 NAb / S-binding IgG for human convalescent panel. Individual animals are shown; bars indicate means.

## Discussion

We previously reported that AS03-adjuvanted CoV2 preS dTM provided robust protection against challenge with SARS-CoV-2 (parental strain USA/WA1-2020, D614) in rhesus macaques and hamsters and provided robust immunity in humans^15, 16^. In this study, we show the CoV2 preS dTM vaccine formulations used as a booster in macaques previously vaccinated with Sanofi Pasteur’s mRNA (D614) or CoV2 preS dTM-AS03 (D614) subunit vaccine candidates induced a robust increase in neutralizing antibody titers against D614G, as early as 7 days post-booster and achieved significantly higher NAb titers than the D35 vaccination peak when adjuvanted with AS03. Importantly, booster injection with CoV2 preS dTM formulations, induced cross-neutralization covering the four currently known SARS-CoV-2 VoC, Alpha, Beta, Gamma and Delta, as well as broadened the NAb response to SARS-CoV-1. The VoC have multiple mutations (Table 1) including E484K, N501Y, K417N and/or L452R in the ACE2 interface of the receptor binding domain (RBD)^19^, some have been associated with increased infectivity (N501Y) and others (E484K) with immune evasion of natural and vaccine-elicited neutralizing antibodies^9, 10^, while SARS-CoV-1 spike protein displays more than 28% of amino acid differences compared to SARS-CoV-2.

**Table 1.**
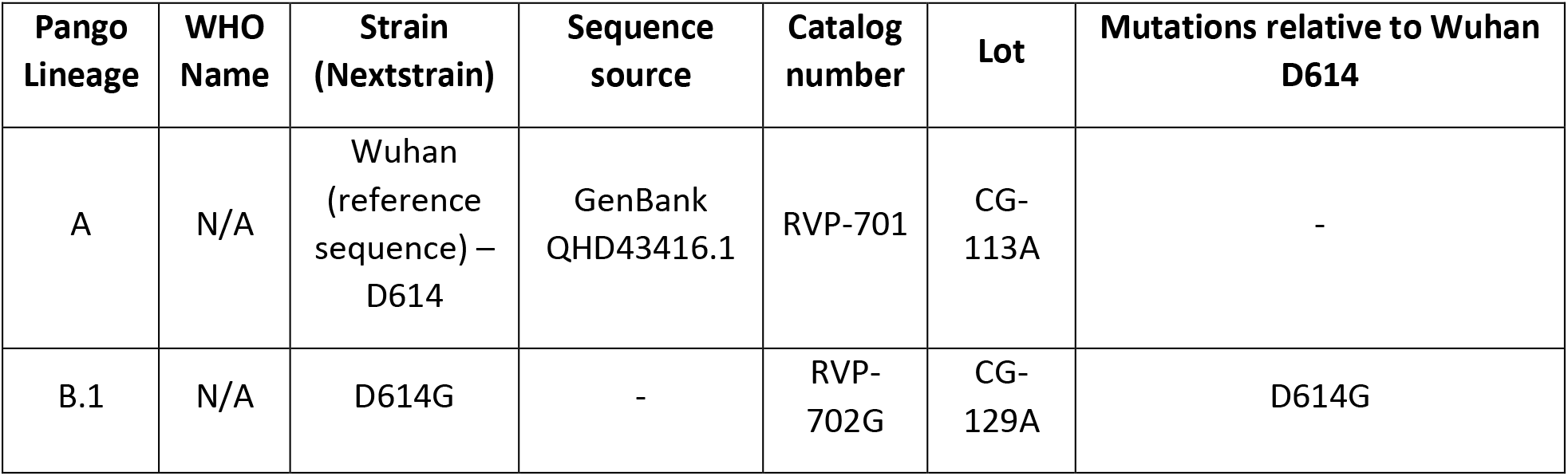
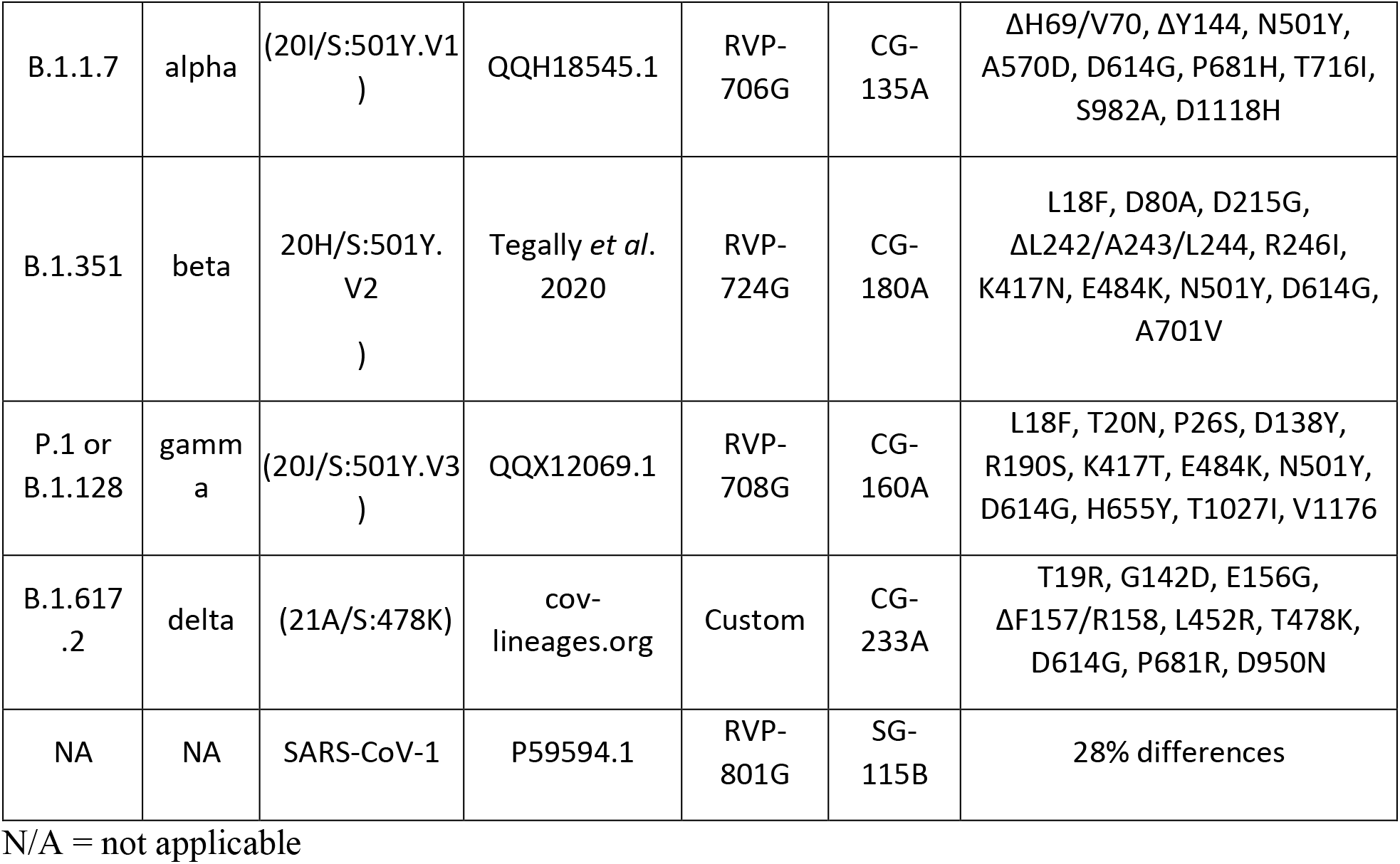

To our knowledge, this is the first report of SARS-CoV-2 booster recombinant vaccine formulations (non-adjuvanted monovalent B.1.351 and AS03-adjuvanted monovalent D614, B.1.351 or bivalent D614+B.1.351) demonstrating broad cross-neutralizing antibodies covering the four current VoC (Alpha, Beta, Gamma and Delta) in macaques primed with different vaccine platforms. Furthermore, in our studies, we observed an extended breadth of neutralization to SARS-CoV-1, which was responsible for the more severe respiratory disease during the 2002-2003 pandemic^20^. Notably, the SARS-CoV-1 spike bares only 72% homology with SARS-CoV-2 spike. It is noteworthy that all booster vaccine formulations tested induced NAb titers with similar breadth, suggesting, in this immunogenicity model, no antigenic sin towards the original vaccine strain was observed. Interestingly, even a booster with the same D614 antigen as used for the primary immunization was able to expand the breadth of neutralization, although there was a trend for a more balanced neutralization profile with the monovalent B.1.351 or bivalent vaccine formulations. Moreover, when used as a booster, no immune interferences were observed in the AS03-adjuvanted bivalent (D614+B.1.351) vaccine formulation.

We observed a significant adjuvant effect of AS03, which increased the titers of neutralizing antibodies against the parental strain, all VoC and SARS-CoV-1 when compared to non-adjuvanted formulation (6 to 8-fold on D614G NAb titers, and 8 to 11.8-fold on B.1.351 NAb titers). This is in line with previous findings in the context of pandemic influenza vaccination, where AS03 adjuvantation consistently resulted in increased magnitude and breadth of the response (including heterotypic antibody responses) in primed individuals^21, 22, 23^.

Interestingly, the non-adjuvanted B.1.351 monovalent formulation was also capable of boosting and broadening the neutralizing responses suggesting that the AS03 adjuvant might not be as necessary for a booster compared to primary vaccination, where it was required to induce functional antibody responses^15^. For comparison, the pseudovirus NAb titers induced by the non-adjuvanted monovalent B.1.351 vaccine were in the same range as those induced in NHP with a third dose of mRNA-1273 vaccine for the Beta, Gamma and Delta variants measured using similar neutralization assays^24^. Based on these encouraging results, non-adjuvanted vaccine candidate and AS03-adjuvanted vaccine formulations using a dose-range of antigen and adjuvant are planned to be further evaluated as booster vaccines in a clinical trial (NCT04762680).

The greater impact of the booster on the neutralization of VoC and SARS-CoV-1 compared to the parental strain (D614) two-dose priming suggests the booster vaccine can remodel the antibody specificity in addition to increasing the antibody levels. Improvement of the antibody quality was also observed in the ratios between functional (neutralizing) and binding antibodies, which increased after the booster immunization. Broader neutralization, coupled to the higher proportion of neutralizing antibody after the booster, is probably due to activation of preexisting memory B cells in primed animals, possibly identifying a key role for memory B cells in mounting recall responses to SARS-CoV-2 antigens and generating cross-reactive NAbs^25^.

While the booster antibody titers reported here cover only the first four weeks post-immunization, the stability of the NAb titers observed after the booster immunization holds promise for an improved durability of the circulating antibodies.

These data have important implications for the utility of current vaccines and inform boosting strategies to address the risks associated with SARS-CoV-2 VoC and waning immunity. As our macaques were not challenged, our study does not define mechanistic correlates of protection against SARS-CoV-2 variants but various studies previously reported that NAbs levels are highly predictive of immune protection from symptomatic SARS-CoV-2 infection^26, 27, 28^. Moreover, a recent report with mRNA vaccine in NHP established the critical role of NAbs as a correlate of protection against infection in NHPs^29^. Based on these studies, and on the levels of NAb titers observed here after the booster against the VoC and SARS-CoV-1, one can hypothesize a booster with CoV2 preS dTM vaccine will augment and extend the protection against existing and newly emerging SARS-CoV-2 variants. In particular, AS03-adjuvanted CoV2 preS dTM booster immunization resulted in higher neutralizing titers than those observed in NHP with a third dose of mRNA-1273 vaccine for the Beta, Gamma and Delta variants^24^, hinting to a beneficial effect of an adjuvanted booster strategy for protection and, potentially, durability of the response.

The study has several limitations. Firstly, the study included a small number of animals, namely 4-5 macaques per group. Secondly, the booster was evaluated in mRNA- and subunit-primed macaques only, as adenovirus vector vaccine primed macaques were not available, however, this priming vaccine is included in the booster vaccine clinical trial (NCT04762680). Thirdly, the interval between the primary vaccination and the booster was 7 months and evaluating shorter and longer intervals would be of interest. Finally, the mechanism underlying the cross-neutralization conferred by the booster, i.e. broadening of the repertoire of antibodies or selection of pre-existing specificities, would need to be further defined. While the antibody repertoire is not easily accessible due to limitation of the immunological tools in macaques, analysis of the Ab quality using system serology and of memory B cell population at later timepoints after the booster immunization are ongoing to better characterize the mechanism for cross-protection.

In conclusion, a single dose of the CoV2 preS dTM recombinant vaccine (non-adjuvanted monovalent B.1.351, AS03-adjuvanted monovalent D614 or B.1.351 or bivalent D614 + B.1.351) used as a booster in macaques previously vaccinated with mRNA or subunit COVID-19 vaccines induced high and stable levels of cross-reactive neutralizing antibodies against the currently known SARS-CoV-2 variants of concerns (Alpha, Beta, Gamma and Delta) and against the two-decade old SARS-CoV-1, as early as 7 days post-boost injection. Our findings show that our vaccine candidates used as a booster have the potential to offer cross-protection against a broad spectrum of variants and have important implications for vaccine control of SARS-CoV-2 VoC.

Clinical studies have been initiated with monovalent (D614 and B.1.351) and bivalent (D614+B.1.351) CoV2 preS dTM vaccine formulations, with or without AS03, to confirm the benefit of a late booster in subjects previously primed with different COVID-19 vaccines including Adenovirus vector-based, mRNA-based and subunit vaccines.

## Materials and methods

### Vaccines

For the primary immunization, the mRNA vaccines were SARS-CoV-2 prefusion Spike constructs 2P, GSAS, 2P/GSAS, 2P/GSAS/ALAYT and 6P/GSAS described in Kalnin *et al*.^18^, the subunit vaccines were AS03-adjuvanted CoV2 preS dTM vaccines, where the antigen were produced using the Phase I/II manufacturing process^16^, 1.3 and 2.6 μg doses, or using an intermediate manufacturing process, 2.4 μg dose.

For the booster, the CoV2 preS dTM derived from the parental D614 strain and the B.1.351 variant was produced using an optimized purification process to ensure a minimum of 90% purity.

The antigens were formulated in monovalent or bivalent formulations adjuvanted with AS03, the monovalent CoV2 preS dTM (B.1.351) were also formulated without AS03. The CoV2 preS dTM was produced from a Sanofi Pasteur proprietary cell culture technology based on the insect cell – baculovirus system, referred to as the Baculovirus Expression Vector System (BEVS). The CoV2 preS dTM (D614) sequence was designed based on the Wuhan YP_009724390.1 strain S sequence, modified with 2 prolines in the S2 region, deletion of the transmembrane region and addition of the T4 foldon trimerization domain. The CoV2 preS dTM (B.1.351) was designed based on the B.1.351 sequence (GISAID Accession EPI_ISL_1048524) and contains the same modifications.

AS03 is a proprietary Adjuvant System composed of α-tocopherol, squalene and polysorbate 80 in an oil-in-water emulsion manufactured by GSK. Vaccine doses were formulated by diluting the appropriate dose of preS dTM with PBS-tween to 250 μL, then mixing with 250 μL AS03, followed by inversion five times for a final volume of 500 μL. Each dose of AS03 contains 11.86 mg α-tocopherol, 10.69 mg squalene and 4.86 mg polysorbate-80 (Tween 80) in PBS.

### Animals and study design

Animal experiments were carried out in compliance with all pertinent US National Institutes of Health regulations and were conducted with approved animal protocols from the Institutional Animal Care and Use Committee (IACUC) at the research facilities. NHP studies were conducted at the University of Louisiana at Lafayette New Iberia Research Center.

Two cohorts of vaccinated NHPs received a booster immunization after randomizing each group within a cohort based on their baseline characteristics (Fig. 1a).

In the mRNA-primed cohort, 16 adult male and female Mauritius cynomolgus macaques (*Macaca fascicularis*) aged 4-10 years, selected based on their responses to the primary vaccination, were randomly allocated to 4 groups of 4 animals according to their baseline characteristics.

In the subunit-primed cohort, 24 adult male Indian rhesus macaques (*Macaca mulatta*) aged 4-7 years were randomly allocated to 5 groups of 4 or 5 animals. In the priming phase, animals received two immunizations of either Sanofi Pasteur’s mRNA COVID (D614) experimental candidate vaccines^18^ or CoV2 preS dTM-AS03 (D614) vaccine through the intramuscular route in the deltoid at day 0 and day 21. Seven months after the primary immunization, both cohorts were immunized with CoV2 preS dTM (B.1.351) without adjuvant, CoV2 preS dTM (D614)-AS03, CoV2 preS dTM (B.1.351)-AS03 and a bivalent CoV2 preS dTM (D614+B.1.351)-AS03, all groups received a total dose of 5 μg of CoV2 preS dTM antigen. An additional group in subunit-primed cohort received a bivalent CoV2 preS dTM (D614+B.1.351)-AS03 high antigen dose (5+5 μg). All immunologic analyses were performed blinded on serum collected at 7, 14, 21 and 28 days post-boost injection. Animal studies were conducted in compliance with all relevant local, state, and federal regulations and were approved by the New Iberia Research Center.

### Convalescent human sera

Convalescent serum panel (N=93) was obtained from commercial vendors (Sanguine Biobank, iSpecimen and PPD). The serum samples were collected within 3 months following PCR positive diagnosis of COVID-19.

The WHO International Standard for anti-SARS-CoV-2 immunoglobulin (human) (NIBSC code: 20/136) was also used.

### Pseudovirus-based virus neutralization assays

Serum samples were diluted 1:4 or 1:20 in media (FluoroBrite phenol red free DMEM + 10% FBS + 10 mM HEPES + 1% PS + 1% Glutamax) and heat inactivated at 56°C for 30 minutes. Further, 2-fold, 11-point, dilution series of the heat inactivated serum were performed in media. Diluted serum samples were mixed with reporter virus particle (RVP)-GFP (Integral Molecular) listed in the table below (Table 1) diluted to contain ~300 infectious particles per well and incubated for 1 hour at 37°C. 96-well plates of ~50% confluent 293T-hsACE2 clonal cells in 75 uL volume were inoculated with 50 uL of the serum + virus mixtures and incubated at 37°C for 72h. At the end of the 72-hour incubation, plates were scanned on a high-content imager and individual GFP expressing cells were counted. The neutralizing antibody titer was reported as the reciprocal of the dilution that reduced the number of virus plaques in the test by 50%.

### Enzyme Linked Immunosorbent Assay (ELISA)

Nunc microwell plates were coated with SARS-CoV S-GCN4 (GeneArt, expressed in Expi 293 cell line) protein at 0.5 ug/ml in PBS overnight at 4°C. Plates were washed 3 times with PBS-Tween 0.1% before blocking with 1% BSA in PBS-Tween 0.1% for 1 hour. Samples were heat inactivated at 56°C for 30 minutes and plated at a 1:450 initial dilution followed by 3-fold, 7-point serial dilutions in blocking buffer. Plates were washed 3 times after a 1-hour incubation at room temperature before adding 50 ul of 1:8000 Rabbit anti-human IgG (Jackson Immuno Reserarch) to each well. Plates were incubated at room temperature for 1hr and washed 3x. Plates were developed using Pierce 1-Step Ultra TMB-ELISA Substrate Solution for 6 minutes and stopped by TMB STOP solution. Plates were read at 450 nm in SpectraMax plate reader. Antibody titers were reported as the highest dilution that is equal to 0.2 OD cutoff.

### Statistical analyses

For both, mRNA-primed and CoV2 preS dTM-AS03-primed cohorts, at the time of the assignment, the characteristics at baseline (sex, age, weight) were balanced in order to have comparable groups. The pseudovirus neutralizing titers were also taken into account as well as the previous vaccine groups. ELISA titers and neutralizing titers were log_10_ transformed prior to statistical analysis. All statistical tests were two-sided, and the nominal level of statistical significance was set to α=5%. The analyses were performed on SEG SAS v9.4^®^.

Statistical comparisons were performed among different groups and conducted as follow for each readout and each strain separately: One way ANOVA with the group as factor were performed to analyze the IgG ratio, PsV ratio post-booster and post prime. Mixed effect models for repeated measures, including group, day, and their interactions, where day was specified as repeated measure, were used for PsV neut / IgG ratio and also to evaluate the boost effect. No adjustment was performed as only 4 to 5 NHPs per groups were available.

## Author Contributions

VL, CB, VP, TT, NA, AR, SG, RC, CG and MK contributed to the concept or design of the study. MK, DH and NA collected and analyzed data. TT provided study coordination. CG, MK contributed critical reagents. NA, CB, VP, VL, TT, AR, SG, RC, CG and MK were involved in the analysis and interpretation of the data. VL, RC provided supervision. VP and CB drafted the first manuscript and all authors critically reviewed manuscript versions. All authors had full access to the data and approved the manuscript before it was submitted by the corresponding authors.

## Acknowledgement

The authors thank Jon Smith for coordinating the production and providing the vaccine antigens for the study, Caroline Ruat for project management, Tong-Ming Fu for coordinating the sharing of mRNA-primed macaques, and Carlos Diaz-Granados, Stephen Savarino and Saranya Sridhar for critical discussions on the study designs and data analysis.

---

**Supplementary Fig. 1.**
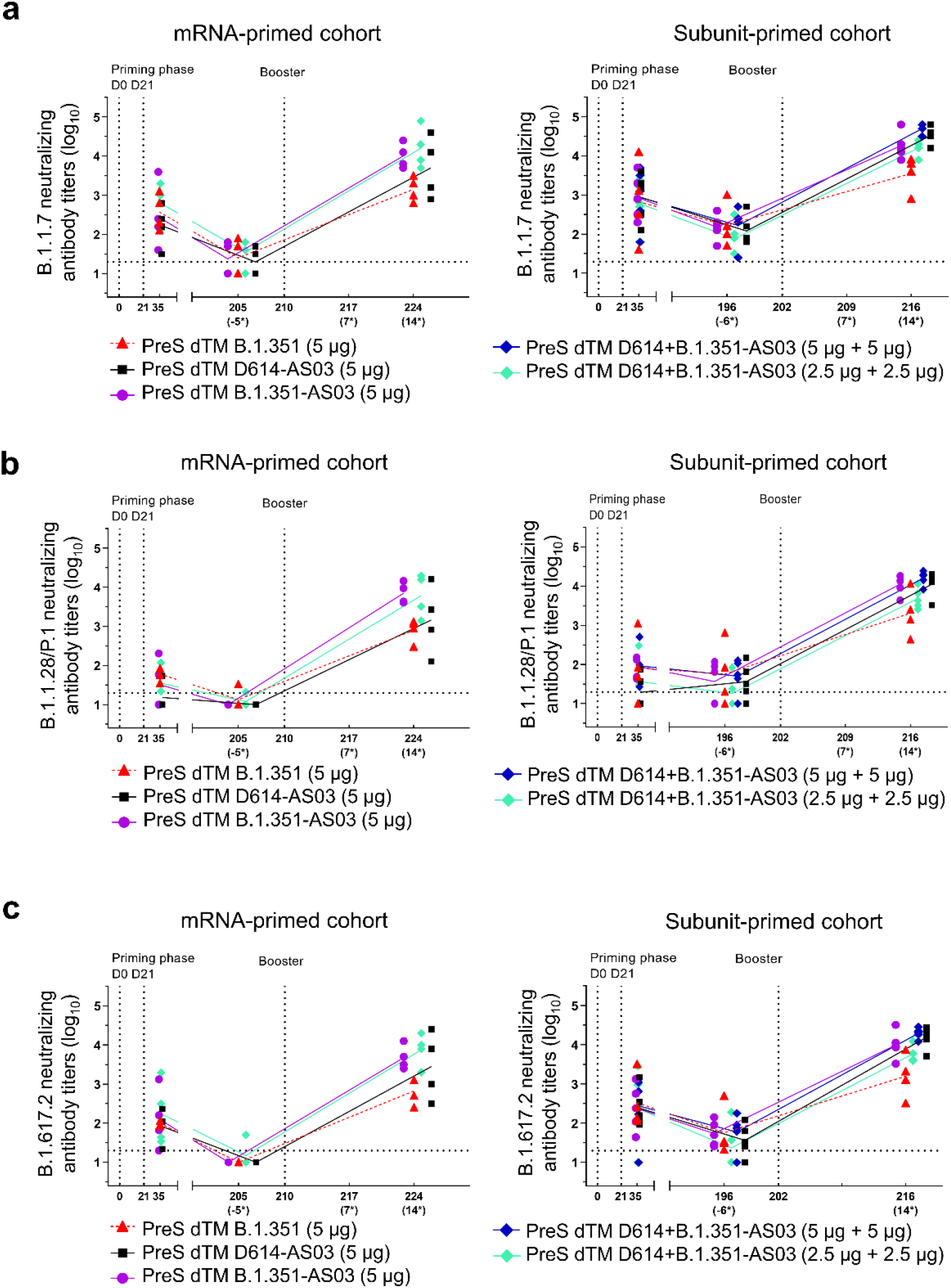
Booster cross-neutralizing antibody responses against Alpha, Gamma and Delta variants in the mRNA- and subunit-primed macaques. Pseudovirus neutralizing antibody (NAb) titers against **a** the SARS-CoV-2 Alpha (B.1.1.7), **b** Gamma (B.1.1.28/P1) and **c** Delta (B.1.617.2) variants were assessed at D35 and 7 months after the priming phase and 2 weeks after the booster immunization in macaques. Individual macaque data are shown.Connecting lines indicate mean responses and horizontal dotted lines the limits of quantification of the assay. Asterisks = timepoints relative to boosters.

**Supplementary Fig. 2.**
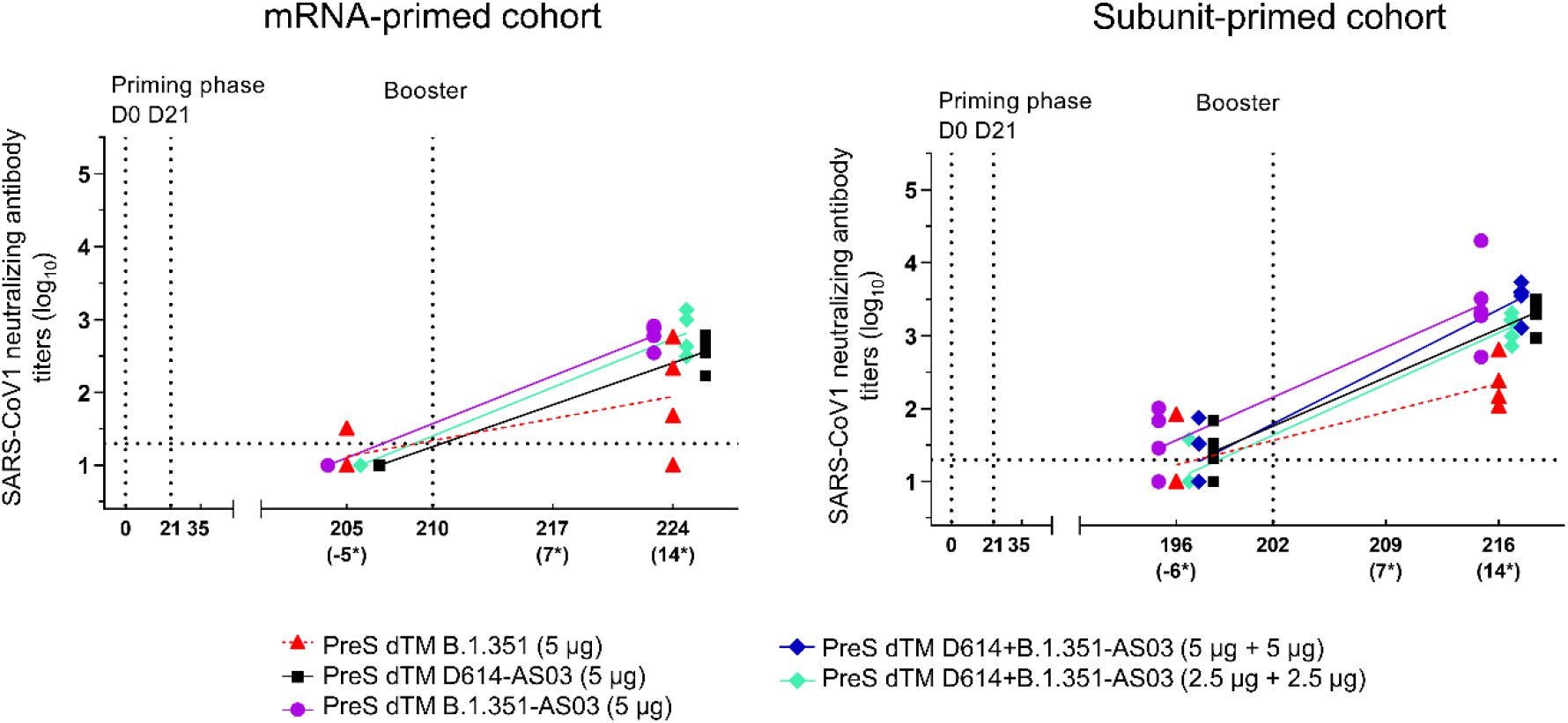
Booster cross-neutralizing antibody responses against SARS-CoV-1 in the mRNA- and subunit-primed macaques. Pseudovirus neutralizing antibody (NAb) titers against SARS-CoV-1 were assessed 7 months after the priming phase and 2 weeks after the booster immunization in macaques. Individual macaque data are shown. Connecting lines indicate mean responses and horizontal dotted lines the limits of quantification of the assay. Asterisks = timepoints relative to boosters.

